# Southeastern Australian Montane Fens Harbour Distinct Microbial Communities Rich in Novel Diversity

**DOI:** 10.64898/2026.05.28.728627

**Authors:** Calum J Walsh, Andrew H Buultjens, Liam KR Sharkey, Louise M Judd, Timothy P Stinear, Sacha J Pidot

**Affiliations:** Department of Microbiology and Immunology at the Doherty Institute, University of Melbourne, Melbourne, 3000, Australia; Centre for Pathogen Genomics, Doherty Institute, University of Melbourne, Melbourne, 3000, Australia

**Keywords:** Montane fens, Microbial ecology, Australian soil, Metabarcoding

## Abstract

Montane fens are rare and microbiologically poorly characterised wetland ecosystems in south-eastern Australia, and their microbial communities remain virtually unexplored. Here, we profile the microbiomes of Victorian montane fens using 16S rRNA metabarcoding of 12 soil cores collected along a 12-m transect and sampled across four depth horizons. Surface soils exhibited slightly higher alpha diversity than deeper layers, but the most pronounced differences occurred in community composition, with surface microbiomes significantly distinct from all subsurface depths. To contextualise these communities within global environmental diversity, we compared them with 9662 Earth Microbiome Project samples spanning 24 environmental materials processed using comparable methods. Montane fen microbiomes were one of the most diverse environmental materials analysed and compositionally distinct from all comparator biomes. Overrepresentation analysis identified signature microbial taxa, including archaeal lineages from the Crenarchaeota and Methanomicrobia and bacterial phyla such as Acidobacteria, highlighting taxa involved in ecological processes associated with acidic, saturated, and organic-rich soils. Notably, the majority of sOTUs detected in montane fens were unique to this environment - the highest proportion of source-specific taxa among all biomes analysed. Together, these findings demonstrate that southeastern Australian montane fens harbour a highly distinctive and largely uncharacterised microbial community, underscoring their ecological uniqueness and the importance of conserving these rare alpine wetlands.

**Data Summary:** Sequencing data is available in SRA BioProject PRJNA1398590; accessions SRR36684598 through SRR36684645. Metadata and accessions for collected montane fen samples are included in Table S1 and metadata for Earth Microbiome Project samples included in this study are listed in Table S4.

**Impact Statement:** Wetland ecosystems are increasingly recognised as important reservoirs of microbial diversity, yet many remain poorly characterised in global microbiome surveys. In this study, we provide the first characterisation of microbial communities inhabiting montane fens in southeastern Australia and place them in a global context using publicly available environmental microbiome data. We show that these fens harbour exceptionally diverse microbial communities that are compositionally distinct from other environmental sources processed using comparable methods, with a high proportion of taxa that are not present in any other sample in an existing reference dataset.

By extending global comparisons to an under-sampled wetland type, this work adds to the growing body of evidence that significant microbial diversity remains undocumented in geographically and ecologically restricted environments. The findings are relevant to researchers working in microbial ecology, environmental genomics, and biogeography, as well as those interested in wetland function and conservation. While largely descriptive, this study represents an important step in expanding environmental genome catalogues and provides a baseline framework for future genomic, functional, and mechanistic investigations of montane wetland microbiomes.

## Introduction

Montane fens on the Mount Baw Baw plateau in the southeastern Australian state of Victoria represent one of the region’s most distinctive and least-studied wetland microbial ecosystems. Occupying gently sloping headwater basins where persistent saturation, cool temperatures, and slow decomposition promote peat formation, these systems form part of a broader complex of alpine and sub-alpine mossbeds distributed across the Victorian Alps^1^. Although often grouped under the umbrella of ‘mossbeds’, Baw Baw’s wetlands are floristically and biogeographically unusual: they lie at the extreme western limit of mainland alpine vegetation and contain several locally endemic or highly restricted plants^2^. Hydrologically, these fens play a critical role in regulating water movement through mountain catchments, retaining moisture and filtering groundwater inputs^3^. Their conservation value is further elevated by their function as habitat for threatened fauna, most notably the FFG- and EPBC-listed Baw Baw Frog (*Philoria frosti*), which is confined to the wet heathlands and peatlands of the region. Unlike most Victorian high-country peatlands, which have experienced multiple fires since 2003, Mount Baw Baw’s fens have remained unburnt since 1939, providing an increasingly rare reference system for understanding long-term ecosystem stability and recovery potential.

Yet, despite their ecological significance, virtually nothing is known about the microbial communities underpinning carbon turnover, and nutrient cycling in these montane fens. Here, we present the first characterisation of soil microbiomes from Baw Baw’s peatlands, providing foundational insights into the microbial diversity and potential functional capacities that support one of Australia’s most important alpine wetland systems.

## Methods

### Sample collection

Samples were collected along a 12m transect. Peatland cores (25mm in diameter) were obtained and were separated into depth intervals of 15 cm onsite. Each interval sample was bagged individually and stored at 4°C during and after transportation to the laboratory and processed for DNA extraction within 24 hours of collection.

### Microbial DNA extraction, sequencing, and analysis

Genomic DNA was extracted and 16S rRNA gene amplicon sequencing was performed following standard protocols from the Earth Microbiome Project^4^ and sequenced on an Illumina Miseq.

Sequence data was processed in *QIIME2*^*5*^. Quality filtering was performed using default cutoffs, Deblur^6^ was used for sub-operational OTU (sOTU) picking and after trimming reads to 150bp, and taxonomic classification was performed using the sklearn-based bayesian classifier^7^ trained on the Greengenes 13_8 16S database^8^. This database was also used at the reference for phylogenetic tree construction using sepp fragment insertion^9^. Prior to calculation of diversity metrics, samples were rarefied to 10,000 features to eliminate the influence of uneven sequencing depth. Principal coordinates analysis (PCoA) of beta diversity metrics was also conducted in *QIIME2*.

Downstream data processing, visualisation, and statistical analyses were performed in *R*^*10*^ (v. 4.5.0). Alpha diversity was compared across depths using linear mixed-effects models implemented in the *lmerTest*^*11*^ package, with Site included as a random effect to account for intra-site dependence. Pairwise contrasts were obtained using the *emmeans*^*12*^ package and tukey adjustment of multiple comparisons. Beta diversity differences were assessed using PERMANOVA (adonis2) in the *vegan*^*13*^ package, with 999 permutations and a restricted permutation design defined using the *permute* package to constrain permutations within each Site. Benjamini & Hochberg correction was used for multiple comparisons. Differentially abundant sOTUs were detected using *LinDA*^*14*^ with parameters prev.filter = 0.1 and mean.abund.filter = 0.001 using a mixed model framework to account for intra-site dependence. Statistical significance was defined as FDR-adjusted p value < 0.05 and absolute log2 fold change > 1. Taxonomic enrichment patterns were analysed using *TaxSEA*^*15*^ with custom taxon groups defined by grouping sOTUs at higher taxonomic ranks (Family, Order, Class, Phylum).

Earth Microbiome Project samples used for comparison had already been subjected to quality control and processing before download – meaning reads were trimmed to 150bp and sOTUs picked. The sOTU sequences were subjected to taxonomic classification and phylogenetic placement as described above. As before, samples were rarefied to 10,000 features before alpha and beta diversity calculations to eliminate the influence of uneven sequencing depth. Diversity metrics were calculated in *Qiime2* as above.

Repeated sampling metadata was not readily available for EMP samples so intra-site dependence was not accounted for during statistical analyses. Shannon diversity was compared between environmental materials using One-Way ANOVA followed by a post-hoc Dunnett’s test in the *DescTools* package (v. 0.99.60)^16^ to compare all EMP environmental materials against the montane fen samples. Beta diversity was calculated using Unweighted Unifrac distance and differences between environmental materials were calculated using PERMANOVA. The PERMANOVA omnibus test was calculated using the *adonis2* function in the *vegan*^*13*^ package and pairwise comparisons were performed using the *RVAideMemoire*^*17*^ package (v. 0.9-83-11). Overrepresentation analysis was performed using the *enricher* function in the *clusterProfiler*^*18*^ package with qvalueCutoff = 0.05. OTUs detected in at least one montane fen sample were used as the query set, with all OTUs detected across the combined montane fen and Earth Microbiome Project dataset used as the background.

## Results

We characterised the microbial communities of these montane fens using 16S rRNA gene metabarcoding of 12 core samples collected at regular intervals along a 12 meter transect, with each core subsampled at four depth strata (0-15 cm, 15-30 cm, 30-45 cm, and 45-60 cm).

Surface soils (0-15 cm) exhibited slightly higher alpha diversity than deeper layers, although this trend was not statistically significant (Fig. 1A). However, community composition at the surface differed significantly from all deeper samples (PERMANOVA, *p* < 0.05; Fig. 1B, Fig. 1C, Table S2). Relative to subsurface depths, surface samples were enriched for 14 sOTUs and depleted for 15 sOTUs (Fig. 1D), although no consistent higher-level taxonomic enrichment patterns were detected (Table S3).

**Figure 1.**
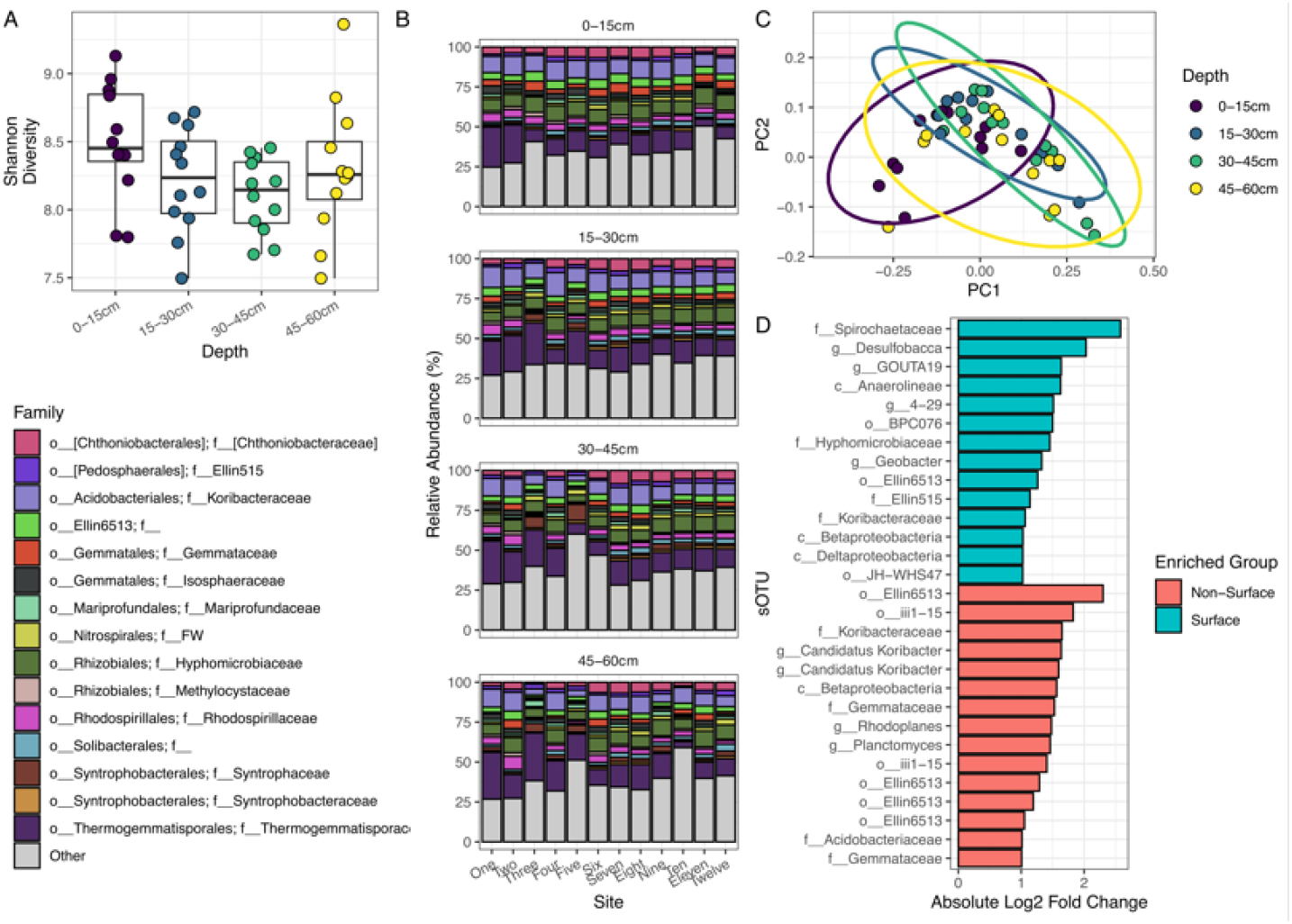
Microbial diversity of Victorian Montane Fens. A) Alpha diversity calculated using the Shannon index. B) Taxonomic profile at the Order level of all samples, stratified by Depth and sampling Site. For readability, only the 15 most abundant families by mean relative abundance are shown, with all others collapsed as “Other”. C) Beta diversity calculated using Unweighted Unifrac distance and ordinated using Principal Coordinates Analysis. Ellipses represent 95% confidence intervals. D) Absolute Log2 Fold Change of sOTUs which are differentially abundant between surface samples and all other depths. Bars are colour filled to show the group in which they are more abundant. Bar labels are the lowest taxonomic rank to which the sOTU could be classified.

To place the montane fen microbiomes in a broader ecological context, we compared them with 9662 Earth Microbiome Project (EMP)^19^ samples representing 24 environmental materials processed using the same protocol (Table S4). Victorian montane fen samples were significantly more diverse than 14 of the 24 comparator environmental materials, only bulk soil and rhizosphere exhibited higher diversity (Dunnet test, *p* < 0.05; Fig. S1, Table S5).

Montane fen communities were compositionally distinct from all other EMP environmental materials, as determined by PERMANOVA analysis of unweighted Unifrac distances (Fig. 2A, Table S6) and overrepresentation analysis (ORA) of sOTU prevalence revealed 23 microbial classes that were significantly overrepresented in montane fen samples relative to the EMP dataset (Fig. 2B). These included archaeal lineages from the phyla Crenarchaeota and Euryarchaeota, as well as several bacterial groups such as Acidobacteria, whose ecological traits provide insight into the biogeochemical processes shaping these wetlands.

**Figure 2.**
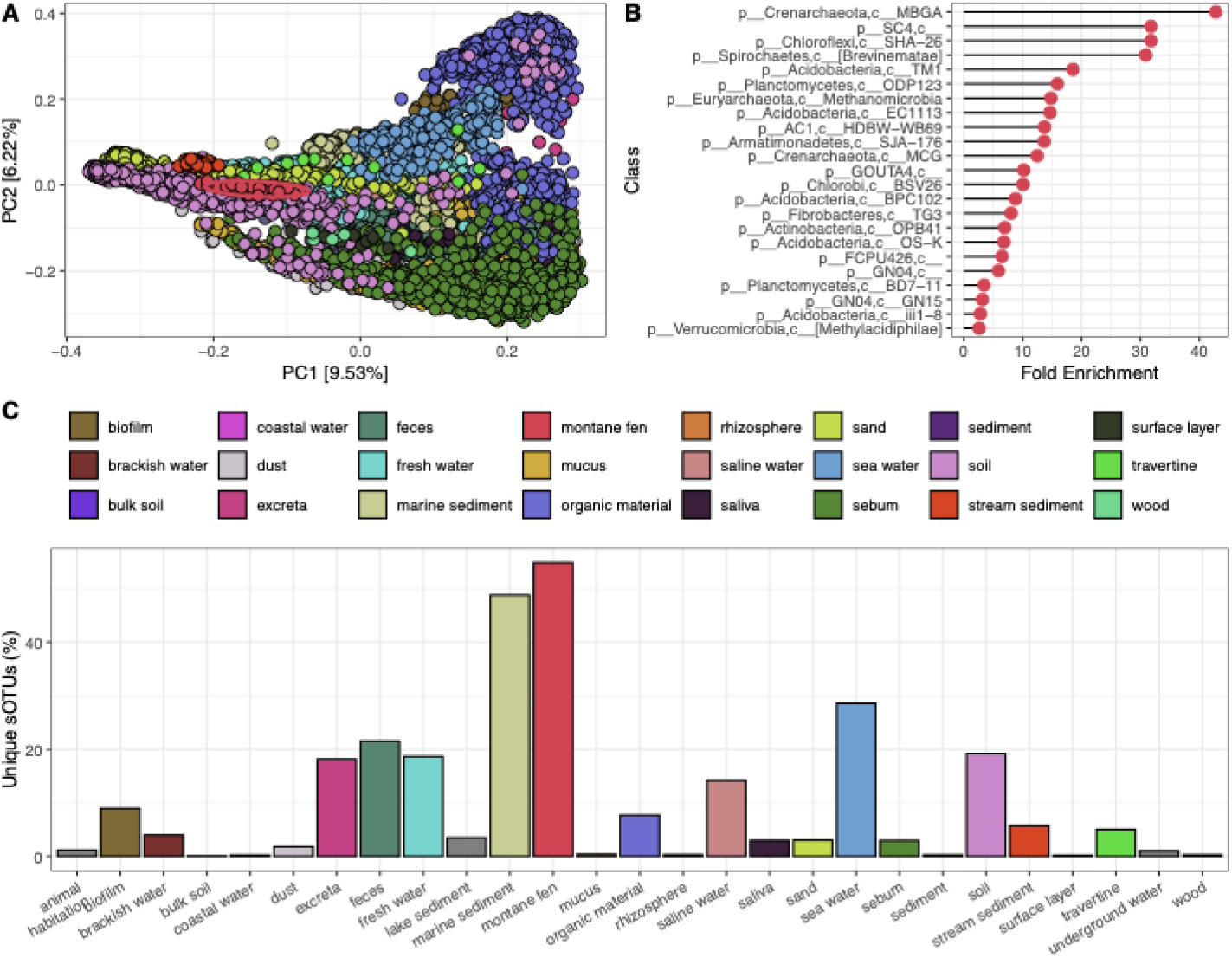
Comparison of Victorian Montane Fen microbiome to 9,662 Earth Microbiome Project samples representing 24 environmental materials. A) Beta diversity calculated using Unweighted Unifrac distance and ordinated using Principal Coordinates Analysis. Montane Fens are highlighted with an ellipse representing the 95% confidence interval. B) Class level analysis of sOTU prevalence showing taxa which are overrepresented in Montane Fens compared to other environmental materials. C) Barchart showing the percentage of sOTUs detected in each environmental material which are unique to that material.

Enrichment of Crenarchaeota, many of which are ammonia-oxidising archaea^20^, suggests an active but tightly constrained nitrification pathway typical of cold, oligotrophic peat systems^21^. This pattern is consistent with slow nitrogen cycling and low decomposition rates that promote peat accumulation. In parallel, the strong representation of Methanomicrobia reflects the anoxic, carbon-rich conditions of water-saturated fen soils, as these are known methane producers^22^, suggesting that montane fens may host microbial networks involved in methane cycling. The overrepresentation of Acidobacteria further aligns with the acidic, organic-rich physicochemical environment characteristic of Australian peatlands^23^. Members of this phylum are often adapted to slow growth under nutrient limitation^24^ and are implicated in the degradation of complex carbon substrates^25^, indicating specialised heterotrophic pathways supporting carbon turnover. Collectively, these patterns indicate that Victorian montane fens harbour a microbiome that is not only compositionally distinct but also selectively enriched for microbial lineages adapted to acidic, water-saturated, and nutrient-limited conditions.

A substantial proportion of the sOTUs detected in montane fen samples (54.8%) were unique to this environment and were not observed in any other EMP sample type (Fig. 2C; Table S5). These fen samples had the highest proportion of source-specific sOTUs among all environmental materials examined, exceeding those observed in marine sediments (48.8%), sea water (28.6%), and faeces (21.5%). Victorian montane fen-specific sOTUs collectively accounted for 40.1 ± 7.8% of total community relative abundance (Fig. S2). Of the 2,581 source-specific sOTUs, 291 were present in at least half of the montane fen samples, and six were detected in all 48 samples (Fig. S3), suggesting the existence of a stable core fen microbiome shaped by long-term hydrological and geochemical conditions.

## Conclusions

Collectively, these taxonomic signatures point to Victorian montane fens harbouring a distinct, redox-structured microbiome shaped by persistent saturation and low nutrient availability. The combination of archaeal nitrifiers, methanogens, and acidophilic heterotrophs underscores the unique biogeochemical niche of montane fens and highlights their ecological importance. The prevalence of taxa that are rare or absent from broader environmental surveys further suggests that montane fens support microbial assemblages that are poorly represented in existing reference collections.

Beyond their distinctive composition, Australian montane fens may function as microbial refugia that promote the persistence and diversification of specialised lineages adapted to these soils. Their restricted geographic distribution and long-term environmental stability may contribute to the maintenance of unique microbial communities with limited dispersal beyond fen environments. As such, these ecosystems offer valuable natural laboratories for investigating peat formation, greenhouse gas dynamics, and the evolutionary processes underpinning microbial adaptation to saturated soils.

By situating montane fen microbiomes within a global comparative framework, this study highlights the importance of incorporating under-sampled ecosystems into large-scale microbial surveys. These findings provide a baseline for future genomic and functional studies aimed at resolving metabolic capabilities, ecological interactions, and climate sensitivity of fen-associated microorganisms, and reinforce the conservation relevance of montane wetlands as reservoirs of microbial biodiversity.

## Author Contributions

Conceptualisation: SJP, TPS. Formal analysis: CJW. Funding acquisition: SJP, TPS. Investigation: SJP, TPS, CJW, AHB, LKRS, LMJ. Resources: SJP, TPS. Supervision: SJP, TPS. Visualisation: CJW. Writing – original draft: CJW. Writing – review & editing: SJP, TPS, LMJ, AHB, LKRS

## Supplementary Material

Supplementary Material file contains supplemental figures and legends for supplemental tables, which are provided as excel files.

