## Supplementary figures and images for "Southeastern Australian Montane Fens Harbour Distinct Microbial Communities Rich in Novel Diversity"

### Supplementary Figure 1

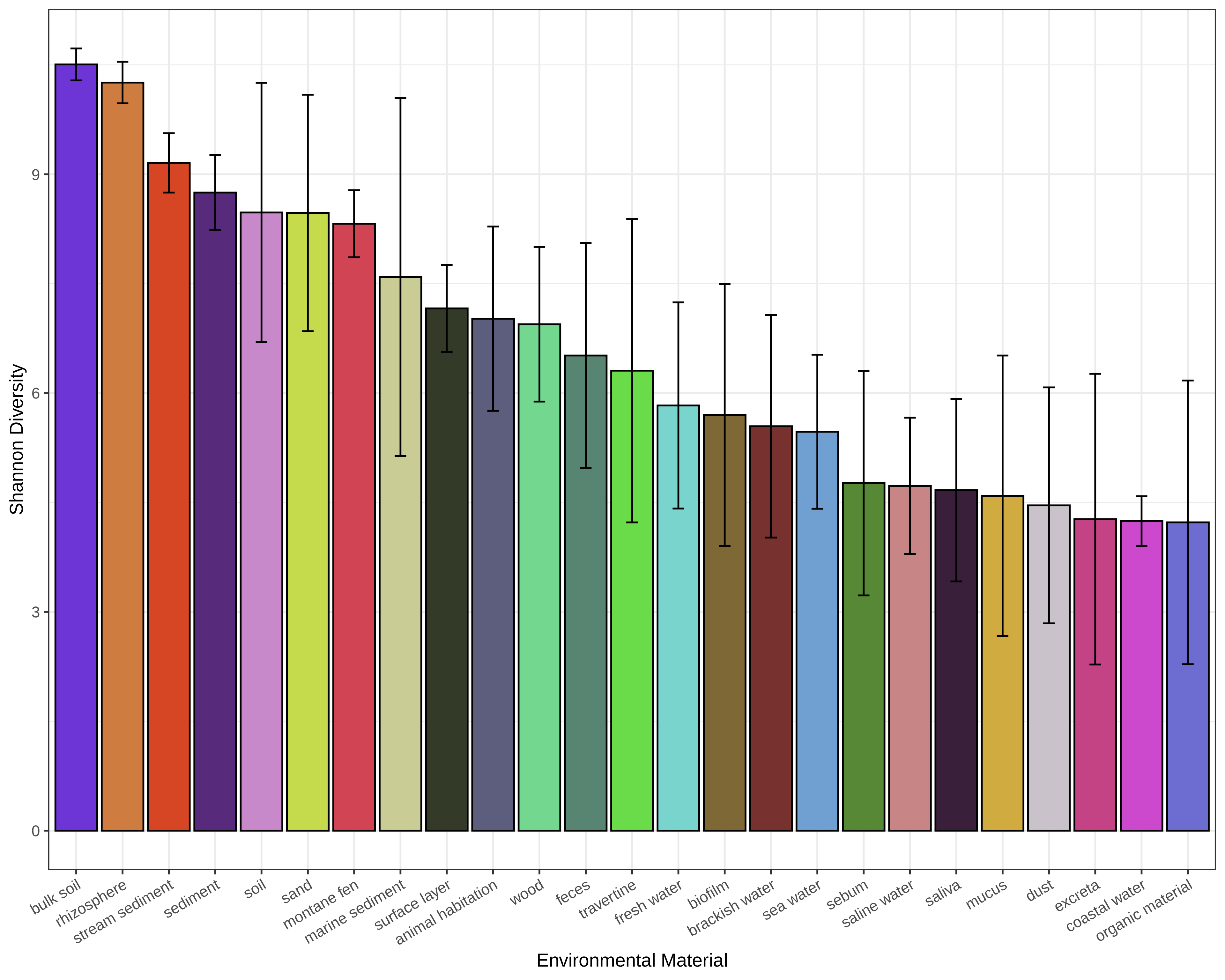

### Supplementary Figure 2

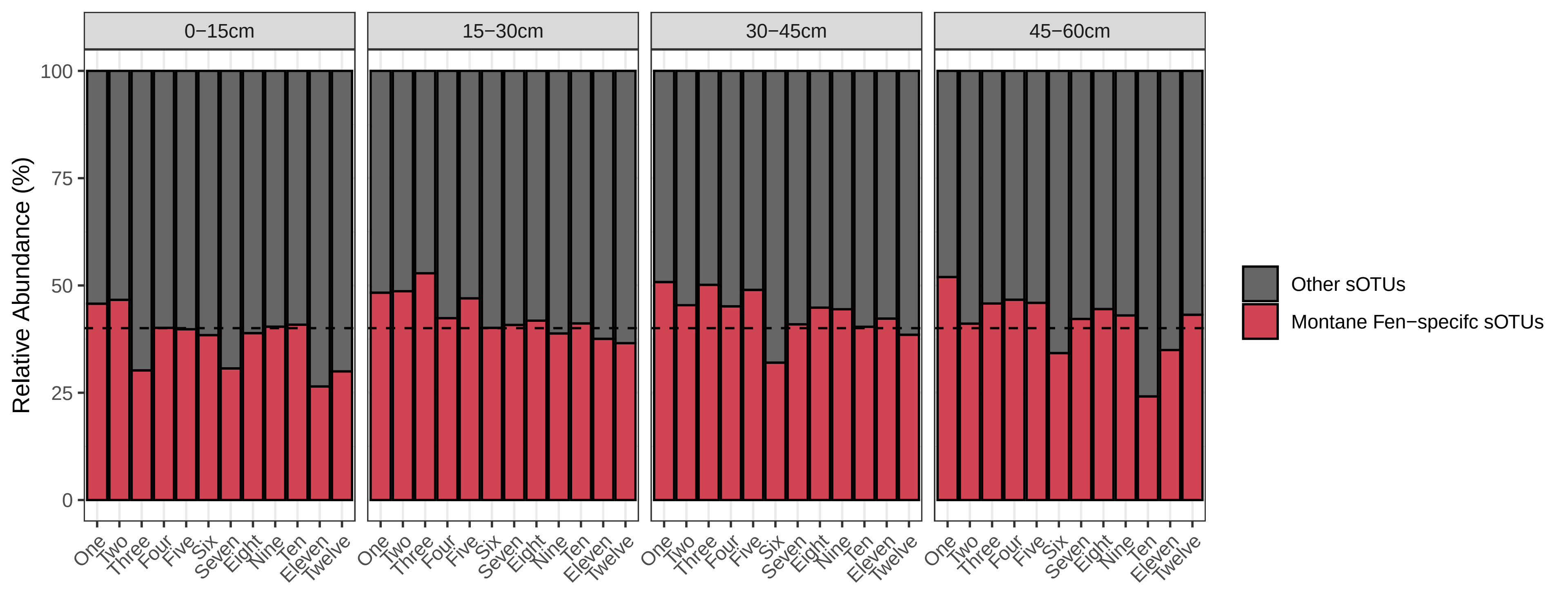

### Supplementary Figure 3

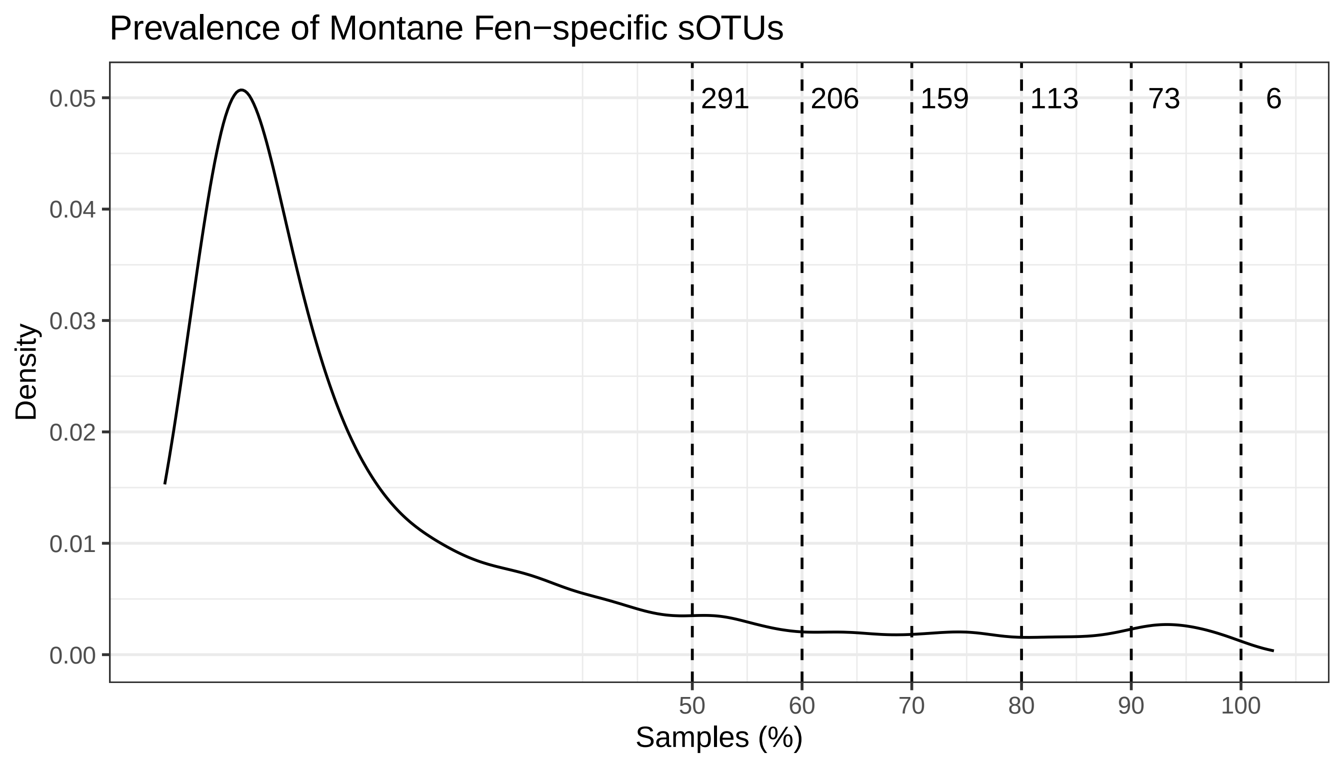
