## Supplementary Material for "Southeastern Australian Montane Fens Harbour Distinct Microbial Communities Rich in Novel Diversity"

**Supplementary Material supporting manuscript "Southeastern Australian Montane Fens Harbour Distinct Microbial Communities Rich in Novel Diversity".**

**Supplemental Figures**

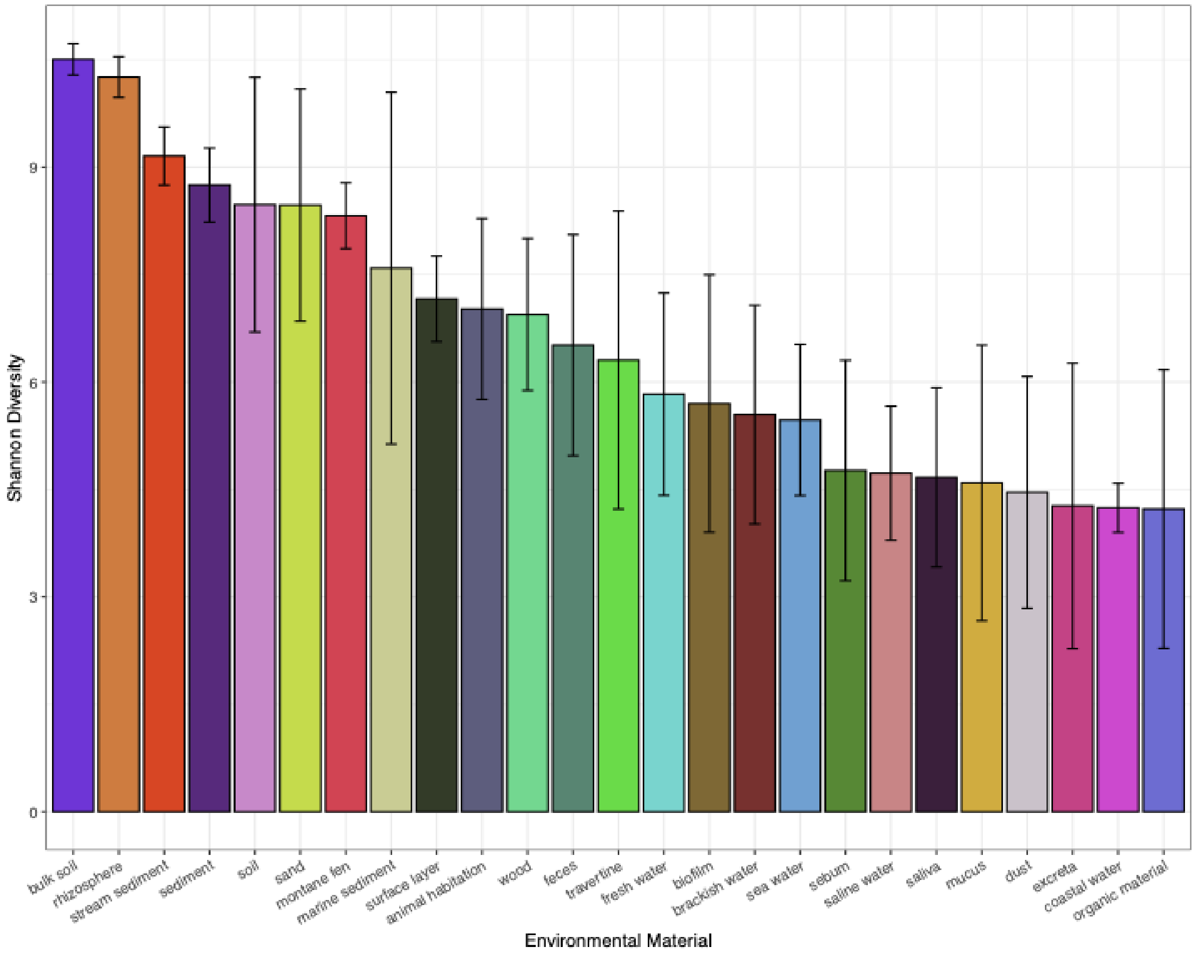


**Figure S1. Microbiome diversity across environmental materials.** Shannon diversity of Victorian montane fen samples compared with other environmental materials from the Earth Microbiome Project. Bars show mean diversity for each material; error bars indicate standard deviation.

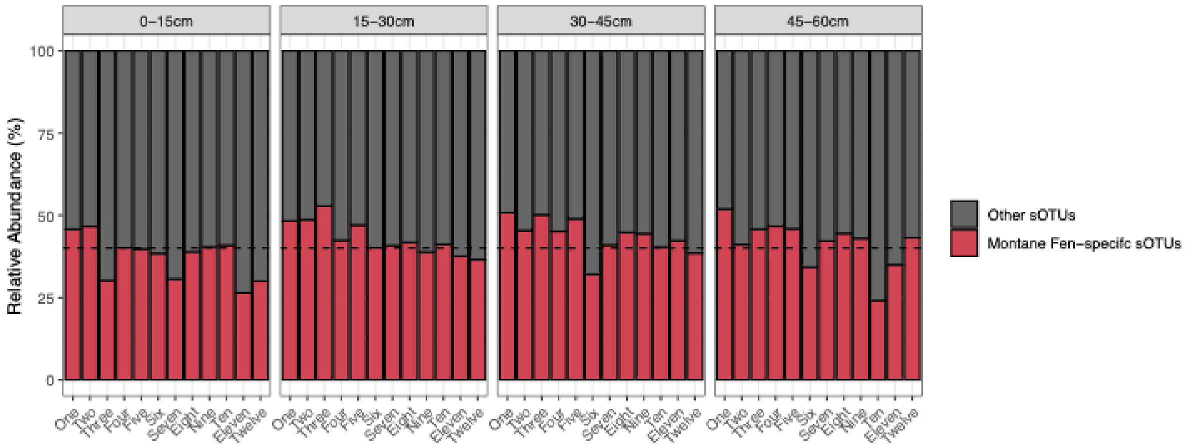


**Figure S2. Relative abundance of montane fen-specific sOTUs across samples.** Per-sample relative abundance of montane fen-specific sOTUs, stratified by depth and site. The dashed line indicates the mean relative abundance of fen-specific sOTUs across all samples (54.8%).

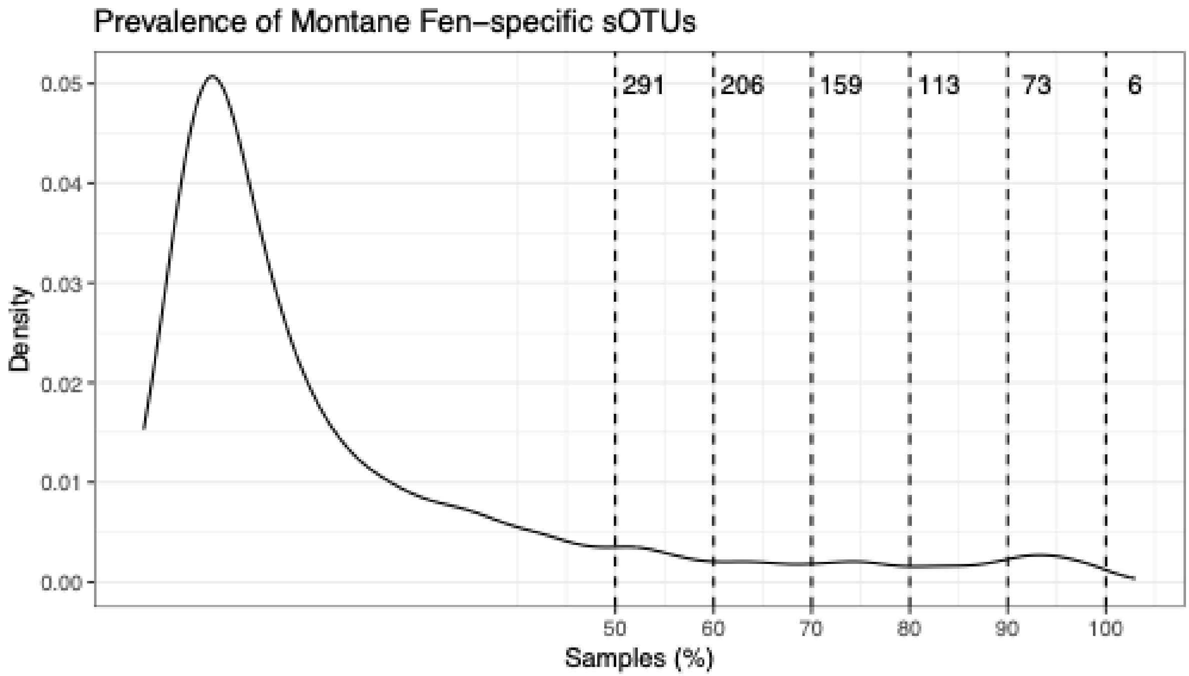


**Figure S3. Prevalence distribution of montane fen-specific sOTUs.** Density distribution of sOTU prevalence, expressed as the percentage of Montane Fen samples in which each fen-specific sOTU was detected. Dashed vertical lines denote core-prevalence thresholds (50-100%), with labels indicating the number of sOTUs present in at least that proportion of samples.

### Supplemental Table Legends

**Table S1. Metadata and SRA accessions of montane fen samples.**

**Table S2. PERMANOVA analysis of montane fen microbiome composition.** Omnibus and pairwise post-hoc PERMANOVA results comparing community composition across sampling depths using four distance metrics. $R^2$ values indicate the proportion of variance explained by depth.

**Table S3. Taxon set enrichment analysis of surface and non-surface samples.** TaxSEA results comparing surface versus non-surface samples using manually defined taxon sets based on higher-rank taxonomy of differentially abundant sOTUs. Analyses were performed at the Family, Order, Class, and Phylum levels.

**Table S4. Sample metadata.** Metadata for all samples included in the comparison between Victorian montane fen samples and the Earth Microbiome Project dataset.

**Table S5. Dunnett's test of alpha diversity across environmental materials.** Post-hoc comparisons of alpha diversity between environmental materials using Dunnett's test, with montane fen samples specified as the reference group.

**Table S6. Pairwise PERMANOVA comparisons between environmental materials.** Pairwise PERMANOVA results assessing differences in microbiome composition among environmental materials using unweighted UniFrac distances. $R^2$ values represent the proportion of variance explained by environmental material.
